# Strain heterogeneity in a non-pathogenic fungus highlights factors contributing to virulence

**DOI:** 10.1101/2024.03.08.583994

**Authors:** David C. Rinker, Thomas J. C. Sauters, Karin Steffen, Adiyantara Gumilang, Huzefa A. Raja, Manuel Rangel-Grimaldo, Camila Figueiredo Pinzan, Patrícia Alves de Castro, Thaila Fernanda dos Reis, Endrews Delbaje, Jos Houbraken, Gustavo H. Goldman, Nicholas H. Oberlies, Antonis Rokas

**Affiliations:** Department of Biological Sciences and Evolutionary Studies Initiative, Vanderbilt University, Nashville, Tennessee, USA; Department of Chemistry and Biochemistry, University of North Carolina at Greensboro, Greensboro, North Carolina, USA; Faculdade de Ciencias Farmacêuticas de Ribeirão Preto, Universidade de São Paulo, São Paulo, Brazil; Food and Indoor Mycology, Westerdijk Fungal Biodiversity Institute, Utrecht, The Netherlands

**Keywords:** comparative genomics, evolution of virulence, fungal pathogenicity, secondary metabolism, specialized metabolism, virulence factor, gliotoxin

## Abstract

Fungal pathogens exhibit extensive strain heterogeneity, including variation in virulence. Whether closely related non-pathogenic species also exhibit strain heterogeneity remains unknown. Here, we comprehensively characterized the pathogenic potentials (i.e., the ability to cause morbidity and mortality) of 16 diverse strains of *Aspergillus fischeri*, a non-pathogenic close relative of the major pathogen *Aspergillus fumigatus*. *In vitro* immune response assays and *in vivo* virulence assays using a mouse model of pulmonary aspergillosis showed that *A. fischeri* strains varied widely in their pathogenic potential. Furthermore, pangenome analyses suggest that *A. fischeri* genomic and phenotypic diversity is even greater. Genomic, transcriptomic, and metabolomic profiling identified several pathways and secondary metabolites associated with variation in virulence. Notably, strain virulence was associated with the simultaneous presence of the secondary metabolites hexadehydroastechrome and gliotoxin. We submit that examining the pathogenic potentials of non-pathogenic close relatives is key for understanding the origins of fungal pathogenicity.

## INTRODUCTION

Filamentous fungi of the genus *Aspergillus* cause a spectrum of diseases collectively known as aspergillosis that account for 340,000 fungal infections every year, and impose a heavy burden on human health and health care systems ^1–4^. The deadliest infections are invasive in nature, with pulmonary aspergillosis being the most common and most deadly type ^5^. Although many species of *Aspergillus* are capable of causing disease, *Aspergillus fumigatus* is the one most frequently seen in the clinic and is considered the primary etiological agent of invasive aspergillosis ^6–11^. Consequently, *A. fumigatus* has been the focus of most studies investigating the mechanisms of fungal virulence in aspergillosis cases.

The genomic, transcriptomic, and metabolic components underlying virulence, and more broadly the pathogenic potential (i.e., the ability to cause morbidity and mortality), of opportunistic fungal pathogens defy ready characterization. Because the mammalian host environment is alien to nearly all fungal species, traits facilitating fungal virulence will be exaptations, that is the coopting of adaptations that originally evolved to serve functions other than virulence ^12, 13^. Traits such as thermotolerance, efficiency and/or flexibility in competing for limited nutritional resources, the ability to grow under hypoxic conditions, as well as the production of an array of secondary metabolites, have all been identified as fungal adaptations to natural challenges that happen to also act as facilitators of opportunistic fungal infections in mammalian hosts ^6, 13^.

Intraspecific diversity can then further broaden the potential of fungi to (accidentally) cause disease, and is especially pronounced in rapidly evolving organisms, such as filamentous fungi ^14^. In *A. fumigatus*, traits such as genotype ^15^, chromatin state ^16^, hypoxia tolerance ^17, 18^, photopigmentation and photoconidiation ^19^, secondary metabolite biosynthesis ^20, 21^, and hypersensitivity to nitrosative or oxidative stresses ^22^, all show extensive variation or heterogeneity between strains. Strain heterogeneity is also a known confounder when predicting clinical outcomes of infection, not only in *A. fumigatus* but in other major fungal pathogens, including *Candida albicans, Nakaseomyces glabrata,* and *Cryptococcus neoformans* ^17, 23–26^.

Strain heterogeneity within pathogenic fungal species can be leveraged to identify key traits contributing to virulence and identify virulence-associated loci ^27^. Comparative analyses between *A. fumigatus* strains have revealed insights into the relationship between hypoxia tolerance and pathogenicity ^17, 26^ and have shown how gene regulatory differences can result in variable responsiveness to fungistatic drugs ^28^.

The relationship between strain heterogeneity and pathogenic potential has been examined in major fungal pathogens but has yet to be examined in non-pathogens. *Aspergillus oerlinghausenensis* and *Aspergillus fischeri* are the two closest known relatives of *A. fumigatus* ^29^; each likely sharing a common ancestor with *A. fumigatus* less than 4 million years ago and both considered to be non-pathogenic ^20^. While only two strains of *A. oerlinghausenensis* are currently known ^30^, *A. fischeri* is globally distributed and can be readily found in the soil, on fruits, on decaying organic matter, and it is a major food spoilage agent ^31–34^. *A. fischeri* is not considered a pathogen and only a handful of aspergillosis cases have been attributed to it ^6, 35–40^. Examining whether strains of non-pathogenic species closely related to major pathogens also vary in their pathogenic potential could inform our understanding of how fungal pathogens originate.

Here we test the hypothesis that strains of a nominally non-pathogenic species exhibit heterogeneity in disease-relevant genes and traits, including in their pathogenic potentials. To this end, we comprehensively characterized the genotypes, chemotypes, and pathogenic phenotypes of 16 *A. fischeri* strains representing different geographic regions and substrates. Using a murine model of pulmonary aspergillosis, we found that *A. fischeri* strains exhibit considerable variation in their virulence profiles and responses to mammalian immune cells. This variation is associated with specific genomic, transcriptomic, and metabolomic differences, suggesting that examination of the pathogenic potential of non-pathogenic close relatives can shed light onto the origins of fungal pathogens.

## RESULTS

### *A. fischeri* strains elicit wide range of responses to murine macrophages

To examine the presence, prevalence, and degree of strain heterogeneity in *A. fischeri*, we selected 16 strains of *A. fischeri* representing a diversity of substrates (soil, food) and geographical origins (Europe, Asia, and Africa; Table 1). We used the *A. fumigatus* CEA17 strain, a clinically derived and highly virulent strain, as a reference ^29, 41^. The growth rate for each strain was measured at both ambient (30°C) and disease-relevant (37°C) temperatures (Methods). All strains grew equally well at 37°C except for one strain (DTO7) and exhibited some variation in pigmented conidiation (Figure S1).

**Table 1:** Strains used in this study.

| VU Strain ID | CBS strain ID | Country of origin | Substrate | Year isolated | Isolated by |
| --- | --- | --- | --- | --- | --- |
| DTO1 | CBS 118456 | Morocco | soil | 2005 | Jos Houbraken |
| DTO2 | CBS 150748 | Thailand | Soil from pineapple field | 2005 | Jos Houbraken |
| DTO3 | CBS 150749 | Spain | Strawberries | 2006 | Jos Houbraken |
| DTO4 | CBS 150750 | Imported in the Netherlands | Heat treated coconut milk | 2006 | Jos Houbraken |
| DTO5 | CBS 150751 | The Netherlands | Cardboard | 2006 | Jos Houbraken |
| DTO6 | CBS 150752 | The Netherlands | Cardboard | 2006 | Jos Houbraken |
| DTO7 | CBS 420.96 | Nepal | soil under *Zea mays* | unkn. | B. v.d. Pol |
| DTO8 | CBS 150753 | Unknown | Soil (glacial) | 2008 | Jos Houbraken |
| DTO9 | CBS 147335 | The Netherlands | Strawberry jam | 2011 | Jos Houbraken |
| DTO10 | CBS 150754 | The Netherlands | Ingredient, maize based | 2011 | Martin Meijer |
| DTO11 | CBS 54465<br>(NRRL181) | Unknown | canned apples | 1923 | C. Wehmer |
| DTO12 | CBS 150755 | Turkey | Soil | 2014 | Rasime Demirel |
| DTO13 | CBS 150756 | South Africa | Soil | 2015 | Martin Meijer |
| DTO14 | CBS 150757 | South Africa | Soil under eucalyptus | 2015 | Martin Meijer |
| DTO15 | CBS 150759 | Belgium | Nickel hydroxide<br>carbonate | 2015 | Martin Meijer |
| DTO16 | CBS 150758 | Italy | Food organic material<br>from biodigester | 2018 | Simone Di Piazza |

We first examined how conidia (asexual spores) from each strain interacted with murine-derived macrophages (Methods). Overall, conidial survival in the presence of macrophages varied widely among strains (Figure 1A). While conidia of some strains had very low survival in the presence of macrophages (e.g., strains DTO2, DTO9, and DTO1), the survivability of conidia from several other strains (e.g., DTO7, DTO3, and DTO16) was high; notably, conidia from strain DTO16 were statistically indistinguishable from conidia from the highly virulent CEA17 reference strain of *A. fumigatus* (unpaired two-sample t-test, Bonferroni-corrected p-value > 0.01).

**Figure 1:**
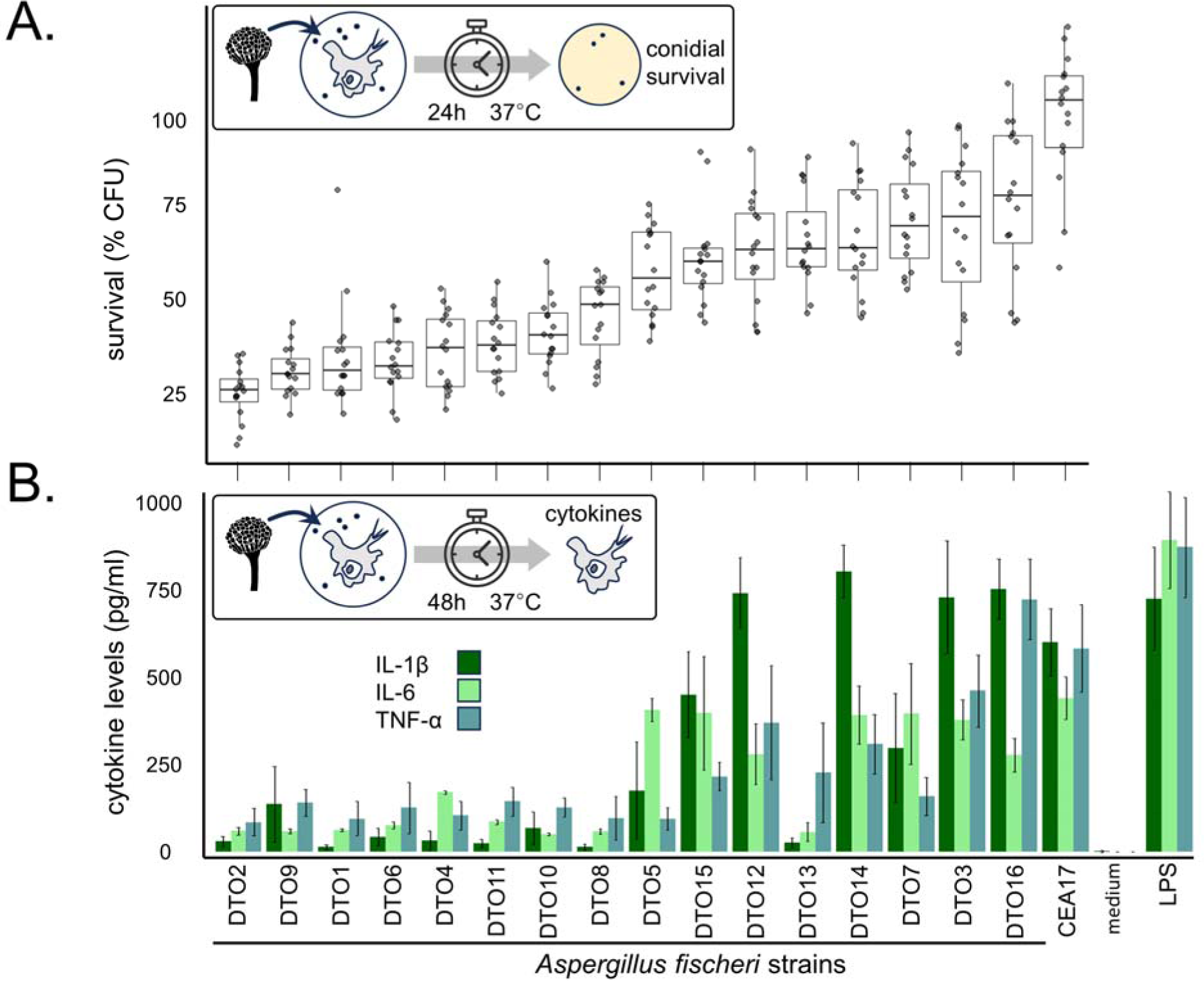
Macrophage response to conidia is highly variable across *A. fischeri* strains. **A**. Survival of conidia (asexual spores) from 16 *A. fischeri* strains (DTO1 - DTO16) as estimated by the number of colony-forming units (CFU) recovered after 24 hours (h) of exposure to bone marrow-derived macrophages (BMDMs) at 37°C. The clinically derived and highly virulent CEA17 strain of the major pathogen *A. fumigatus* was used as a reference. Boxplots show the mean and interquartile range; whiskers show max/min values of data that are within ±1.5× the interquartile range, respectively. **B**. Pro-inflammatory cytokine levels produced by BMDMs after 48h incubation with conidia at 37°C. Lipopolysaccharide (LPS) and cell medium were used as positive and negative controls, respectively. Bars represent the mean of eight separate measurements carried out across two replicates. Error bars show standard deviation.

We next measured the pro-inflammatory cytokine responses of macrophages, an *in vitro* proxy for the mammalian innate immune response, to conidia from each strain (Methods). We found a graded, strain-dependent response in macrophage cytokine levels (Figure 1B). Specifically, some of the cytokine responses elicited by certain strains (e.g., DTO12, DTO14, DTO3, and DTO16), rivaled the responses elicited by the *A. fumigatus* CEA17 strain. Indeed, certain *A. fischeri* strains elicited levels of IL-1β and/or TNF-α that surpassed those elicited by the *A. fumigatus* CEA17 strain (e.g., DTO3, DTO12, DTO14, DTO16); these differences were not statistically significant (all unpaired two-sample t-tests, false discovery rate-corrected p-value > 0.01).

Overall, there was a general concordance between strains that were more resistant to macrophages and those inducing higher levels of proinflammatory cytokines (Figure 1). Notably, there was a large difference in both assays between the macrophage response to conidia from the *A. fumigatus* reference strain CEA17, and those from the *A. fischer*i reference strain NRRL181 (DTO11). Taken together, the conidial challenge assays of 16 strains demonstrate that there exists substantial phenotypic variation within *A. fischeri* that is not reliably reflected by the *A. fischeri* reference strain. Importantly, some levels of conspecific variation are at least on par with levels of interspecific variation.

### *A. fischeri* strains vary considerably in their virulence profiles

The strain heterogeneity of *A. fischeri* observed in the macrophage assays raised the hypothesis that these strains could vary in their pathogenic potential. To test this, we measured the virulence (mortality) of *A. fischeri* strains in a chemotherapeutic murine model of pulmonary aspergillosis (Methods).

We observed substantial variation in virulence across the strains (Figure 2). While no strain of *A. fischeri* exhibited the 100% mortality rate of *A. fumigatus* CEA17, several *A. fischeri* strains (e.g., DTO14, DTO15, DTO12, and DTO16) consistently produced lethality rates of 50% or greater, and four strains (e.g., DTO14, DTO7, DTO9, and DTO8) resulted in earlier onset of death compared to CEA17. Overall, DTO14 was the deadliest strain, resulting in two deaths after only two days and having the lowest animal survival rate of any *A. fischeri* strain (35% mean survival rate over the two trials). Importantly, there was a large difference in the survival of mice exposed to conidia of the *A. fumigatus* reference strain CEA17 compared to those infected by the *A. fischer*i reference strain NRRL181 (DTO11). This difference in lethality between the two reference strains for each species is consistent with previous results ^29^.

**Figure 2:**
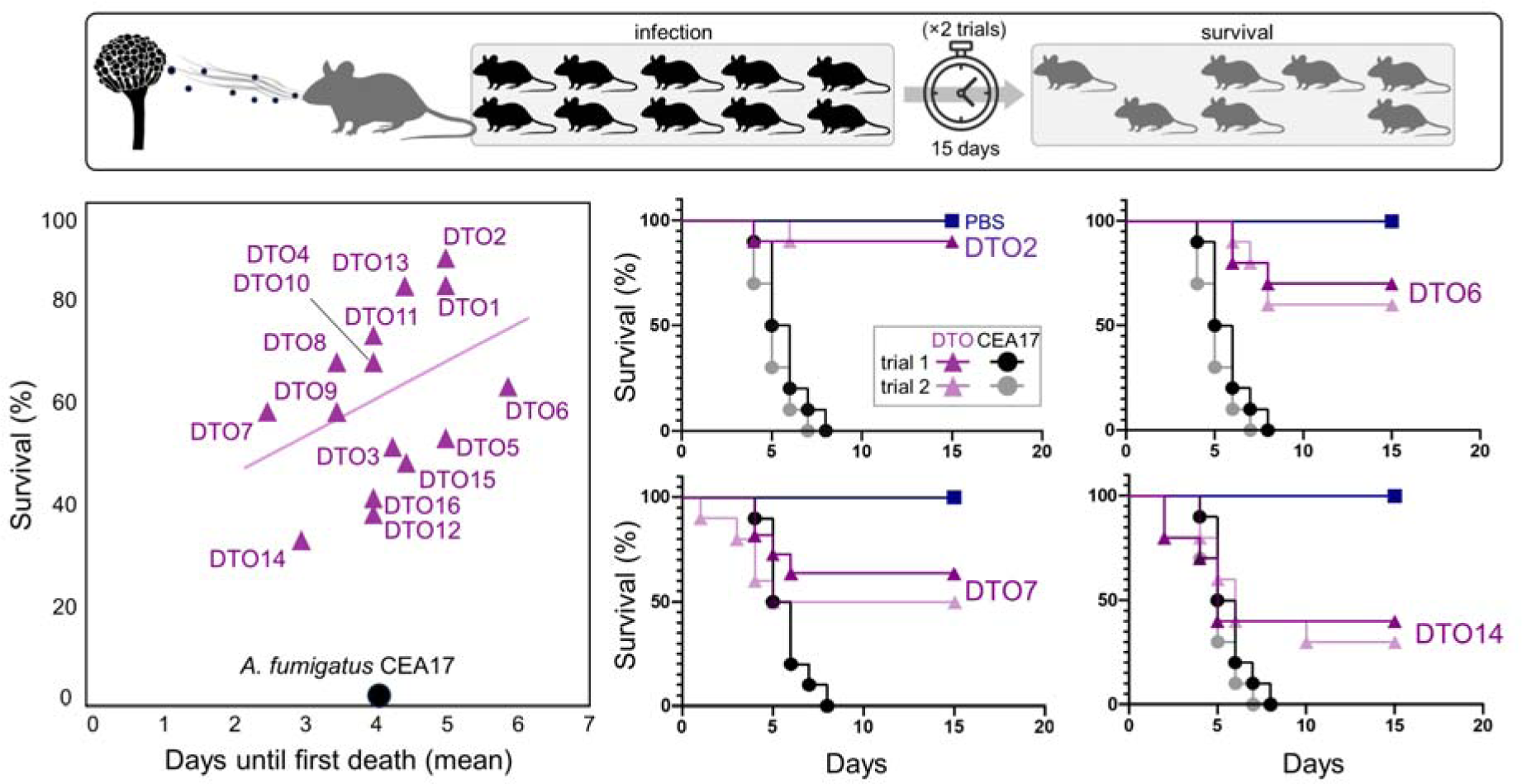
Murine model of pulmonary aspergillosis reveals wide range of virulence across *A. fischeri* strains. Shown are the 15-day survival rates of mice exposed to 16 strains of *A. fischeri*. The clinically derived and highly virulent CEA17 strain of the major pathogen *A. fumigatus* was used as a reference. **Left**: Scatter plot comparing the mean number of days until the first recorded death (x-axis) and the cumulative survival rate (y-axis) for each of the strains. **Right**: Kaplan-Meier curves of the survival rates (y-axis: percent survival, and x-axis: days) for two weakly virulent (DTO2 and DTO6) and two strongly virulent (DTO7 and DTO14) strains of *A. fischeri*.

### Genes associated with virulence and secondary metabolism are highly conserved across *A. fischeri* strains

We next quantified the genetic relationships between these strains and investigated the genomic context of the observed phenotypic diversity. In particular, we evaluated each strain for the presence of 207 genes previously associated with virulence in *A. fumigatus* ^29, 42, 43^. We also assessed intraspecific variation in biosynthetic gene cluster (BGC) content, given the known role played by secondary metabolites in virulence ^44, 45^. To this end, we constructed chromosome-level *de novo* genome assemblies and built gene models for all 16 strains (Methods).

To evaluate evolutionary relationships between the 16 *A. fischeri* strains, we inferred a strain phylogeny using 3,354 single copy Eurotiomycetes (BUSCOs) orthologs present in all strains (Figure 3A; Methods). We found that *A. fischeri* strains are genetically very similar, with the highest pairwise distance occurring between DTO2 and DTO10 (∼4×10^-3^ nucleotide substitutions/site). This is on a par with the distance between the two *A. fumigatus* strains (Af293 and CEA10, which is a progenitor strain of CEA17 ^41^) included in our analysis (∼2.4×10^-3^ nucleotide substitutions/site). In contrast, any *A. fumigatu*s outgroup strain is over 36 times more diverged (∼0.14 substitutions/site) from any *A. fischeri* strain.

**Figure 3:**
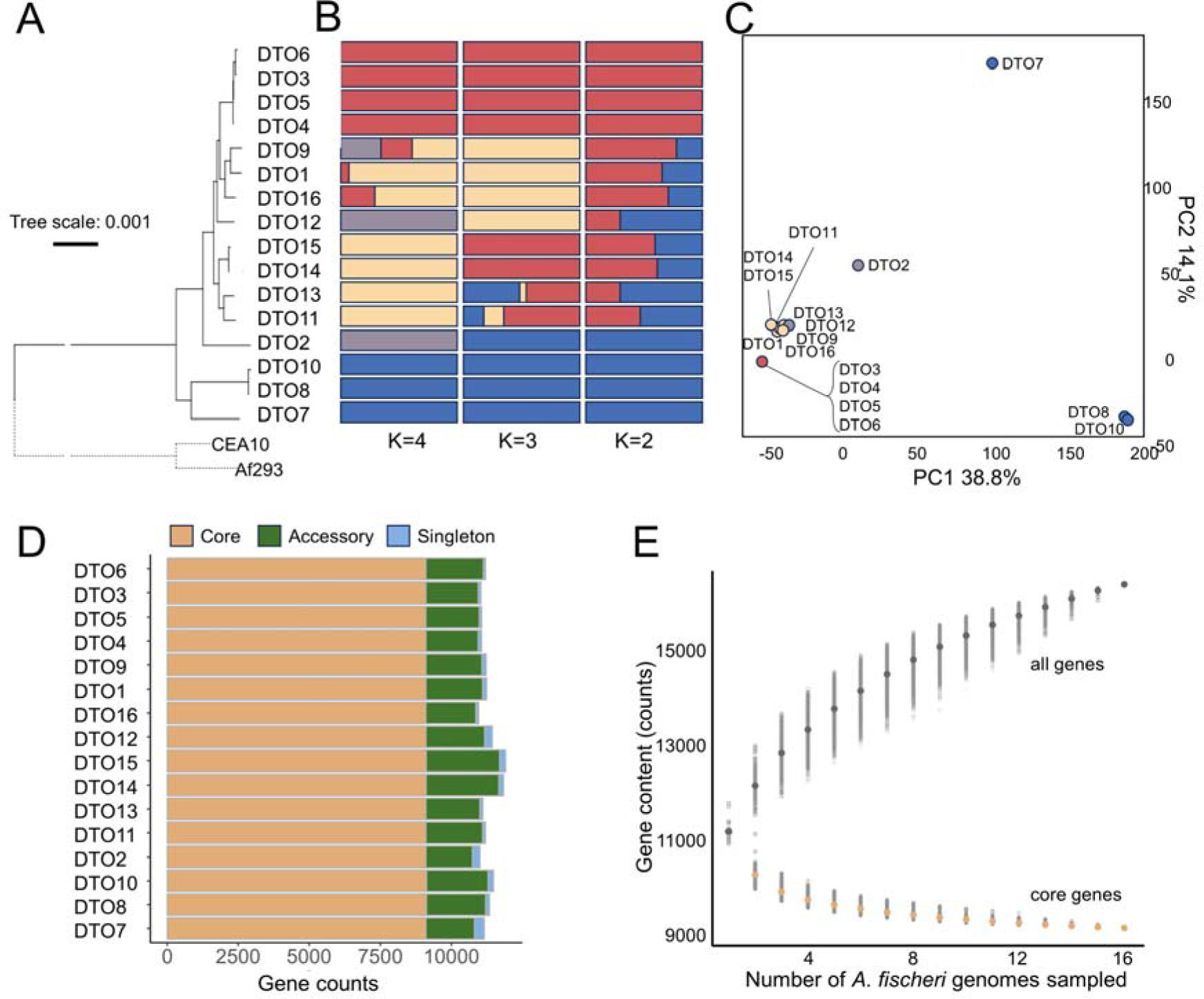
*A. fischeri* comprises at least two populations and has an open pangenome. **A**. Phylogeny of the 16 strains of *A. fischeri* inferred from 3,546 single-copy BUSCO orthologs present in all strains; strains CEA10 and Af293 of *A. fumigatus* were used as outgroups. Bootstrap values were 100 for all branches and are not shown. **B**. Results of ADMIXTURE population inference analyses of 8,763 linkage disequilibrium-pruned single nucleotide variants (SNVs). The optimal number of populations K is either 2 or 3 depending on the parameters chosen. **C.** Relatedness of *A. fischeri* strains as determined by SNV-based discriminant analysis of principal components (DAPC). Colors represent ADMIXTURE groups when the number of populations K=4. **D**. Summary of pangenome results highlighting accessory genes present in two to 15 strains (4,669) and singleton genes (2,426); the *A. fischeri* pangenome also contains 9,053 core genes. Only the longest isoforms are being counted. **E**. Results of incremental, random subsampling of the genomes of 16 *A. fischeri* strains, evaluating each subsample for shared gene content. Subsample size was incrementally increased from one to 16, with 10,000 replicates performed at each step, illustrating that the *A. fischeri* pangenome is open (Heaps’ law regression, γ= 0.858).

To gain insights into the population structure of *A. fischeri*, we performed ADMIXTURE analysis of 215,627 single nucleotide variants (SNVs) that are polymorphic within *A. fischeri* (Methods). These analyses suggested that *A. fischeri* strains can be divided into between 2 and 4 ancestral populations (Figure 3B). This structure is broadly reflective of and consistent with the *A. fischeri* phylogeny (Figure 3A). Further analysis of these SNVs by both discriminant analysis of principal components (DAPC) and principal components analysis (PCA) (Figure 3C and Figure S2, respectively; Methods) consistently identified strains DTO10 and DTO8, as well as DTO2 and DTO7, to be more distinct from the rest, potentially representing populations whose breadth has not been fully captured by our sampling. That said, except for DTO7, these four strains are not notably distinct in terms of their conidial viabilities, cytokine responses, virulence profiles, and overall pathogenic potentials (Figure 2).

Genes from all 16 strains were consolidated into an *A. fumigatus* pangenome comprising 16,148 single-copy pangenes (Methods). Of the pangenes, 9,053 are present in every strain (core genes), with the remaining 7,095 in only some (accessory genes) or one (singleton genes) strain (Figure 3D). The accessory gene component of the pangenome is monotonically increasing without converging, a finding indicative of an open pangenome (Figure 3D; Heap’s law γ =0.85849). Therefore, genome sequencing of additional *A. fischeri* strains will likely reveal more accessory genes.

Among a set of 207 virulence-associated genes previously identified in *A. fumigatus* ^46^, orthologs of all but *pesL* (*Afu5g12730*), *fmaG* (*Afu8g00510*), and *fma-TC* (*Afu8g00520*) are present in most, if not all, *A. fischeri* strains. Even the strain that lacks the highest number of virulence-associated genes (DTO6) still contains 201 of them. Our pangenome-based approach provides further support for the inference of Mead et al. that *A. fischeri* possesses most of the virulence-associated genes identified in *A. fumigatus* ^29^.

Many of the 207 virulence-associated genes are located within BGCs, reflecting the fact that secondary metabolites are important to fungal lifestyles, including pathogenic lifestyles ^13, 44^. We examined six previously described *Aspergillus* BGCs that are responsible for the biosynthesis of seven secondary metabolites variously implicated as virulence factors in *A. fumigatus*: gliotoxin, fumitremorgin A/B, verruculogen, trypacidin, pseurotin, fumagillin, and hexadehydroastechrome (Table 2) ^20, 47^. All the genes for the gliotoxin and hexadehydroastechrome BGCs are present in all strains, while the genes for the trypacidin and verruculogen/fumitremorgin A/fumitremorgin B BGCs are present in all but one and two strains, respectively. The pseurotin and fumagillin BGCs were incomplete (i.e., only some of the genes were present) in all strains. Our expanded taxon sampling highlights the low variability of BGC presence/absence between strains, and augments previous work showing high conservation of BGCs in strains of *Aspergillus* species, including the major pathogen *A. fumigatus* ^21, 48^ and the rare pathogen *A. nidulans* ^49^. Overall, these data show that most genomic features associated with virulence in *A. fumigatus* are present in and conserved among *A. fischeri* strains; in turn, this conservation suggests that pathogenic potential cannot be readily attributed to differences in gene content.

**Table 2:**
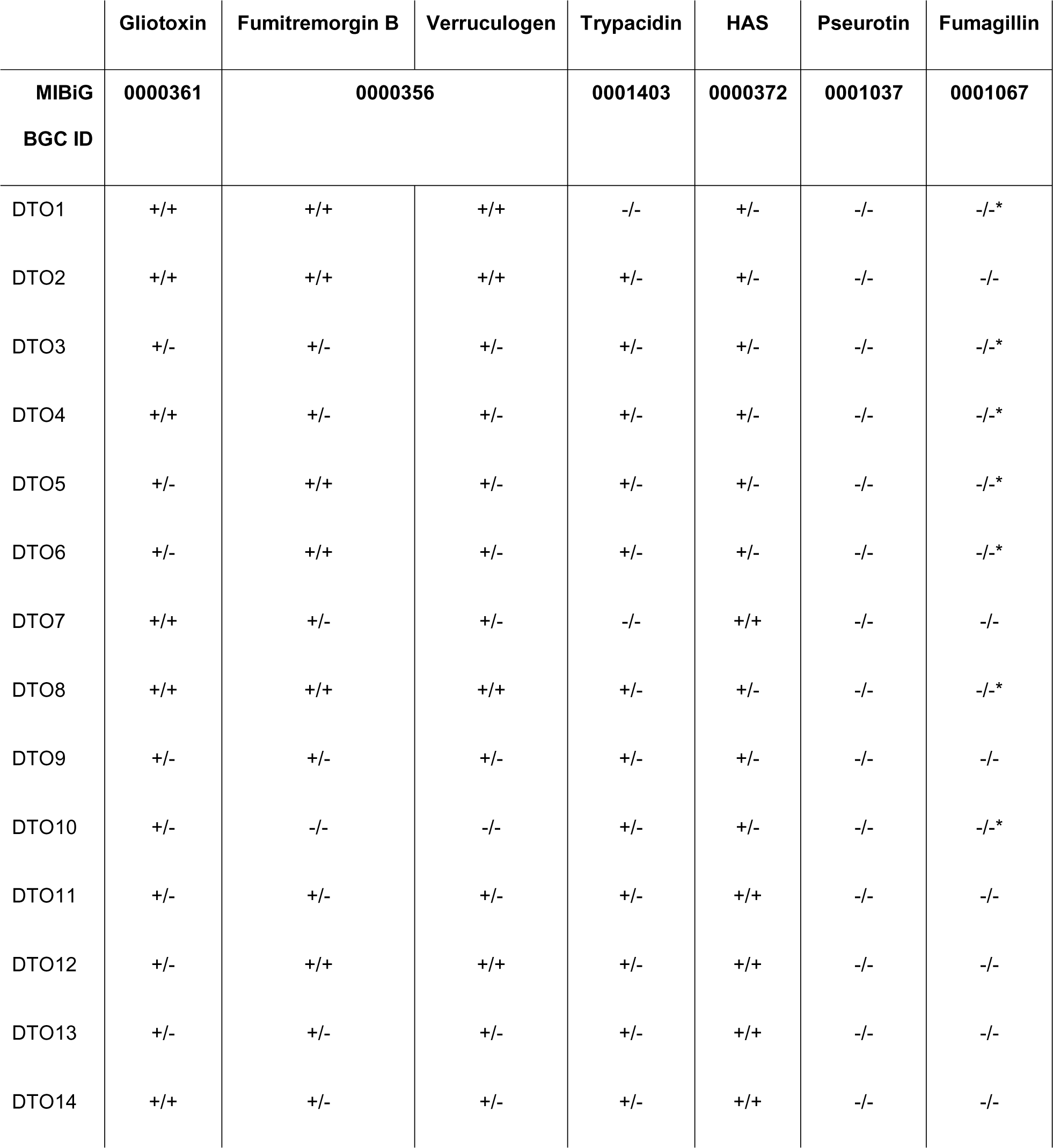
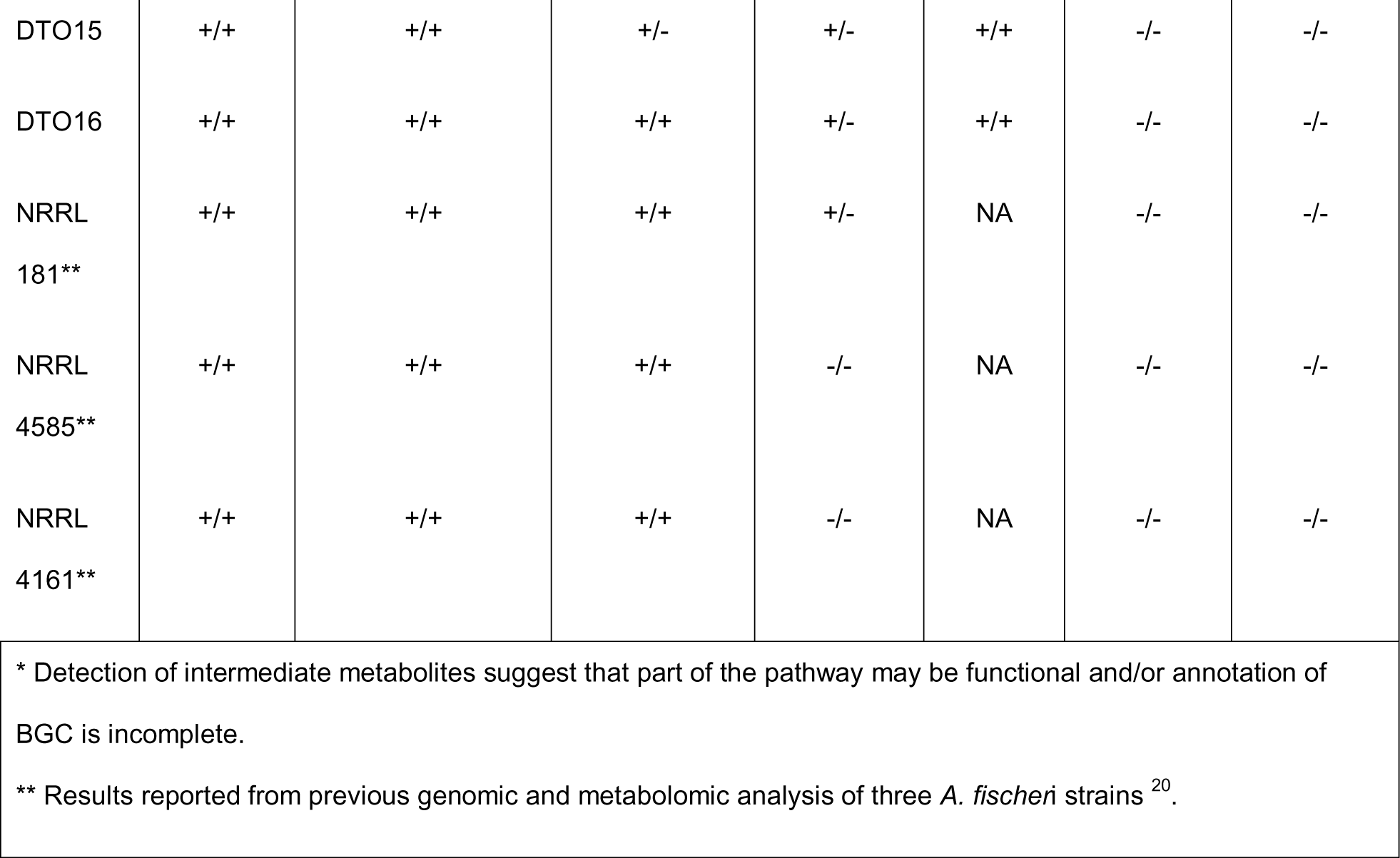
Variation in the presence/absence of select BGCs and their secondary metabolite compounds. Presence (+) and absence or lack of detection (-) of BGCs and their compounds indicated as (BGC/compound).

|  | Gliotoxin | Fumitremorgin B | Verruculogen | Trypacidin | HAS | Pseurotin | Fumagillin |
| --- | --- | --- | --- | --- | --- | --- | --- |
| MIBiG<br>BGC ID | 0000361 | 0000356 |  | 0001403 | 0000372 | 0001037 | 0001067 |
| DTO1 | +/+ | +/+ | +/+ | -/- | +/- | -/- | -/-* |
| DTO2 | +/+ | +/+ | +/+ | +/- | +/- | -/- | -/- |
| DTO3 | +/- | +/- | +/- | +/- | +/- | -/- | -/-* |
| DTO4 | +/+ | +/- | +/- | +/- | +/- | -/- | -/-* |
| DTO5 | +/- | +/+ | +/- | +/- | +/- | -/- | -/-* |
| DTO6 | +/- | +/+ | +/- | +/- | +/- | -/- | -/-* |
| DTO7 | +/+ | +/- | +/- | -/- | +/+ | -/- | -/- |
| DTO8 | +/+ | +/+ | +/+ | +/- | +/- | -/- | -/-* |
| DTO9 | +/- | +/- | +/- | +/- | +/- | -/- | -/- |
| DTO10 | +/- | -/- | -/- | +/- | +/- | -/- | -/-* |
| DTO11 | +/- | +/- | +/- | +/- | +/+ | -/- | -/- |
| DTO12 | +/- | +/+ | +/+ | +/- | +/+ | -/- | -/- |
| DTO13 | +/- | +/- | +/- | +/- | +/+ | -/- | -/- |
| DTO14 | +/+ | +/- | +/- | +/- | +/+ | -/- | -/- |
| DTO15 | +/+ | +/+ | +/- | +/- | +/+ | -/- | -/- |
| DTO16 | +/+ | +/+ | +/+ | +/- | +/+ | -/- | -/- |
| NRRL<br>181** | +/+ | +/+ | +/+ | +/- | NA | -/- | -/- |
| NRRL<br>4585** | +/+ | +/+ | +/+ | -/- | NA | -/- | -/- |
| NRRL<br>4161** | +/+ | +/+ | +/+ | -/- | NA | -/- | -/- |

\* Detection of intermediate metabolites suggest that part of the pathway may be functional and/or annotation of BGC is incomplete.
\*\* Results reported from previous genomic and metabolomic analysis of three *A. fischeri* strains<sup>20</sup>.

### *A. fischeri* strains show high variation in transcriptional response to elevated temperatures

The pathogenicity of opportunistic fungi is a result of both their ability to thrive within the host environment and the incidental effects of their responses (e.g., secondary metabolite production) to growth under disease-relevant conditions ^13, 44, 50, 51^. To characterize genes that may be active, or may become active, in infection-relevant conditions, we profiled the transcriptomes of each of the strains at 30°C and 37°C (Methods).

Strains showed very diverse transcriptional profiles in response to growth at 37°C. Strain-to-strain comparisons of differential expression profiles do not follow the intraspecific relationships inferred from the strain phylogeny (Figure 4A). For example, the expression profile of strain DTO16 is most similar to those of DTO15 and DTO14, but the strain is not the sister group to the DTO14+DTO15 clade on the phylogeny (Figure 4A). This is not surprising given that transcriptional variation is not expected to always mirror strain evolutionary history.

**Figure 4:**
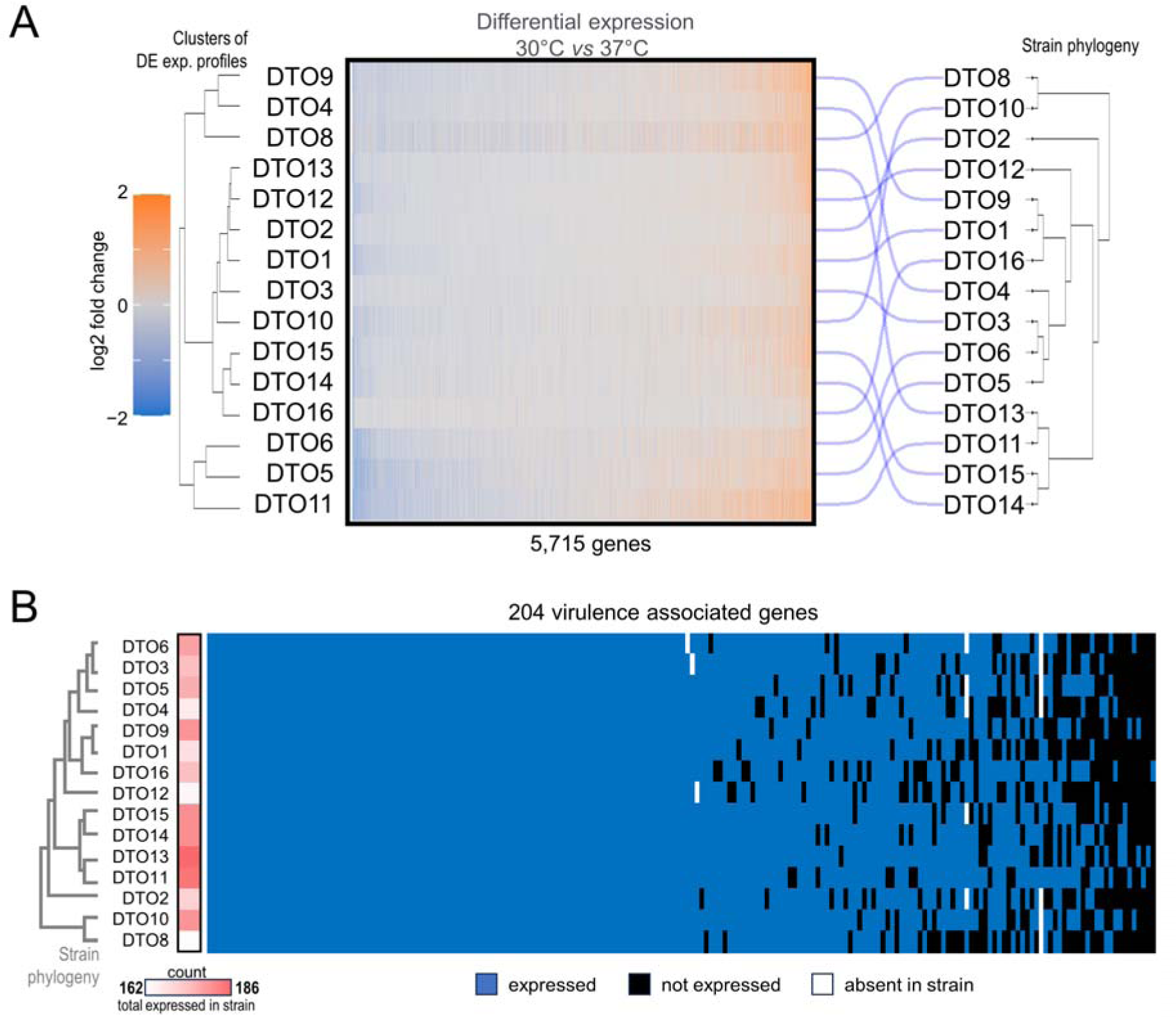
Strain heterogeneity in the transcriptional response to the disease-relevant condition of growth at 37°C. **A**. Temperature-dependent (30°C vs 37°C) differential expression of 5,715 genes (x-axis) in each of 15 strains of *A. fischeri* (y-axis). The differential expression profiles of strains are clustered hierarchically (left) and their clustering is compared to the strain phylogeny using a tanglegram (right). Strains showed very diverse transcriptional profiles in response to growth at elevated temperatures. Heatmap intensities reflect the log2 fold changes of each gene between the two treatment conditions. **B.** Distribution of 204 virulence-associated genes (x-axis) in each of 15 *A. fischeri* strains (y-axis); the strain phylogeny is shown at the far left. Virulence associated genes are shown as being present and expressed (blue), present and not expressed (black), or not present. Heatmap (red) reflects the number of virulence-associated genes present in each strain. All 204 genes were expressed in at least one strain, and more than half (108) were expressed in all strains. Among the strains, DTO13 showed the most (185) virulence-associated genes expressed and DTO8 the least (162). Note: strain DTO7 was excluded from transcriptome profiling because of poor growth at 37°C.

Examining this temperature-dependent transcriptional response with respect to virulence reveals that the three least virulent strains (DTO1, DTO2, and DTO13; Figure 2) and the three most virulent strains (DTO14, DTO15, and DTO16; Figure 2) group together with three other strains (DTO3, DTO10, and DTO12) based on transcriptional profiles (Figure 4A). Conversely, strains that have identical phenotypes in mouse and macrophage assays are genetically and transcriptionally distinct (DTO10 and DTO4).

No genes are uniquely expressed in either the most- or the least-virulent strains, and only two genes (orthologs of *Afu5g13940* and *Afu6g07260*) are exclusively upregulated in the most virulent strains. While the functional annotations of both genes suggest that they could be involved in secondary metabolite biosynthesis and export (oxidoreductase activity and nucleobase transmembrane transporter, respectively), neither is predicted to be located within a BGC.

Of the 204 virulence-associated genes identified in *A. fischeri*, transcripts for 162 (79%) are detected in all strains, with some strains expressing as many as 186 genes (91%) of the full complement (Figure 4B). In contrast with the global differential expression profiles, the profiles of virulence-associated genes more closely followed the *A. fischeri* phylogeny (Figure 4B). While there appeared some occasional relationship between the presence/absence of virulence-associated gene expression and virulence in mice (e.g., the highly virulent DTO14, DTO15, and DTO16 strains ranked within the top quartile for number virulence-associated genes expressed, while the weakly virulent DTO1 and DTO2 strains were in the bottom quartile), there were plenty of exceptions (e.g., the weakly virulent DTO13 had the highest number of virulence-associated genes expressed, while the second-to-lowest number of virulence-associated genes were expressed in the highly virulent DTO12).

Expression of genes within the four virulence-associated BGCs present in *A. fischeri* (Table 2) showed pervasive transcriptional activity in most of the strains (Figure 5). Genes in the verruculogen/fumitremorgin A/fumitremorgin B BGC displayed the largest positive changes in their levels of expression in response to the elevated temperature, with DTO9 and DTO12 showing positive regulation across the entirety of the BGC. Genes of the hexadehydroastechrome BGC were also broadly upregulated. Conversely, genes in the trypacidin BGC showed the lowest levels of gene expression; multiple genes were not expressed across the strains and no single strain showed expression for all gene members of the trypacidin BGC. Interestingly, the remaining genes of the otherwise incomplete fumagillin BGC were active and upregulated at 37°C (Figure S3).

**Figure 5:**
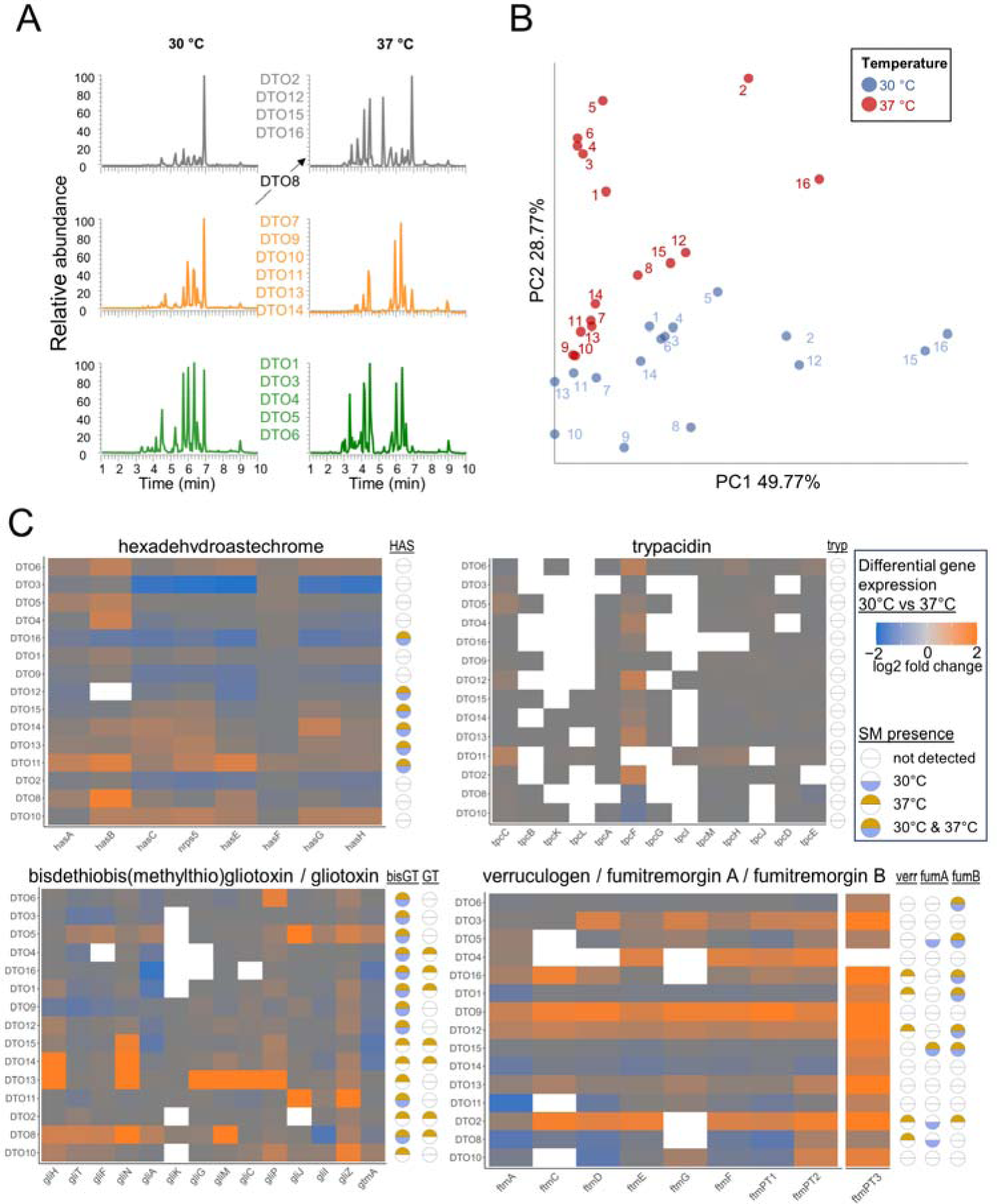
Variation in secondary metabolite production among strains is principally driven by strain diversity. **A.** Chromatograms of the 16 strains of *A. fischeri* displayed three distinct profiles at both 30°C and 37°C temperatures. Shown are representative chromatograms of each profile. The chromatograms report retention times (x-axis) and signal intensities normalized to the highest value (y-axis) of produced metabolites; different peaks correspond to different metabolites. **B**. Principal component analysis based on retention times and individual peak areas for different metabolites from each of the 16 strains following growth at either 30°C or 37°C. Approximately 50% of the variation in the chemistry profiles comes from variation between strains (PC1), and 29% of the variation is due to temperature (PC2). **C**. Temperature-dependent differential expression of transcripts for constituent genes in each of four, virulence-associated biosynthetic gene clusters in 15 *A. fischeri* strains (DTO7 was excluded because of poor growth at 37°C). Secondary metabolite (SM) presence/absence at 30°C and 37°C is indicated to the right of each heat map. White cells on the heatmap indicate lack of transcriptional activity. HAS: hexadehydroastechrome; tryp: trypacidin; bisGT: bisdethiobis(methylthio)gliotoxin; GT: gliotoxin; verr: verruculogen; fumA: fumitremorgin A; and fumB: fumitremorgin B.

### Variation in secondary metabolite production across strains corresponds to variation in virulence

To evaluate the secondary metabolite profiles of *A. fischeri* strains at 30°C and 37°C, we assayed extracts from cultures of each of the strains by ultraperformance liquid chromatography–mass spectrometry (Figure 5A; Methods). Overall, 30% of the chemodiversity of secondary metabolite profiles was due to growth temperatures (30°C or 37°C) while nearly 50% was attributable to strain heterogeneity (Figure 5B). Some of the most metabolically distinct strains included several of the most- (DTO12, DTO15, DTO16) and least- (DTO1 and DTO2) virulent strains.

We next examined the presence of seven virulence-associated secondary metabolites previously identified in *A. fumigatus* and whose BGCs were also characterized above ^20, 52^. Four of these virulence-associated secondary metabolites (gliotoxin, hexadehydroastechrome, fumitremorgin A/B, and verruculogen) appeared very frequently or ubiquitously among our strains, while three other compounds (fumagillin, pseurotin, and trypacidin) were not detected.

Examination of the secondary metabolites detected against the virulence profiles across *A. fischeri* strains revealed noteworthy patterns in gliotoxin and hexadehydroastechrome. Gliotoxin was observed in seven strains and its inactive derivative bis(methylthio)gliotoxin was present in all strains (Figure 5B). In the context of infection, gliotoxin acts to suppress the innate immune response in mammalian hosts ^53^. *A. fischeri* strains with detectable levels of gliotoxin at 37°C correspond to those showing elevated virulence (greater than 50% mortality) in mice. However, the presence of gliotoxin was not in itself sufficient to predict elevated virulence as gliotoxin was also present in DTO2, the least virulent strain (Figure S4).

Another secondary metabolite detected in only a subset of *A. fischeri* strains was hexadehydroastechrome, an intracellular iron chelator that has been associated with increased virulence in *A. fumigatus* ^47, 52, 54^. In *A. fischeri*, nearly every gene of the hexadehydroastechrome BGC was transcriptionally active in all strains, but the compound was only detected in six strains (Figure 5). Among those six strains, four (DTO12, DTO14, DTO15, and DTO16) showed elevated virulence in mice.

Notably, we detected both hexadehydroastechrome and gliotoxin at 37°C in 3 of the 4 most virulent strains, making their co-occurrence a statistically significant chemical predictor of virulence in a mouse model of pulmonary aspergillosis (binary logistic regression χ2=7.9913, p value=0.0047; Figure 5C; Figure S4). We hypothesize that this association may be meaningful to the elevated pathogenic potential exhibited by these strains and may be mechanistically explained by the complementary roles of gliotoxin and hexadehydroastechrome in confounding host defenses, while simultaneously protecting the pathogen. Specifically, gliotoxin blunts active host immune response, and has recently been shown to induce ferroptosis in mammalian cells. Ferroptosis is a form of cell death triggered by aberrantly elevated levels of intracellular iron and can affect macrophages directly^55–57^. Fungi are also susceptible to ferroptosis ^58^ and the iron chelating property of hexadehydroastechrome could act to protect fungi from a similar fate by sequestering excess iron. None of the other secondary metabolites we detected among our *A. fischeri* strains exhibited this degree of correspondence between their presence/absence (either alone or in combination), and virulence.

## DISCUSSION

The comprehensive characterization of the chemotype, genotype, and pathogenic phenotype of 16 strains of the non-pathogen *A. fischeri* revealed pervasive strain heterogeneity. Of the traits measured, one of the most surprising findings was the range of virulence displayed across strains. In a murine model of pulmonary aspergillosis, mortality ranged from just 10% for the least virulent strain, to infections that killed up to 70% for the most virulent strain. Such variation in lethality between strains of a non-pathogenic species complicates not only how we distinguish pathogenic from non-pathogenic fungal species, but also how specific traits subtending that virulence are disentangled (Figure 6).

**Figure 6:**
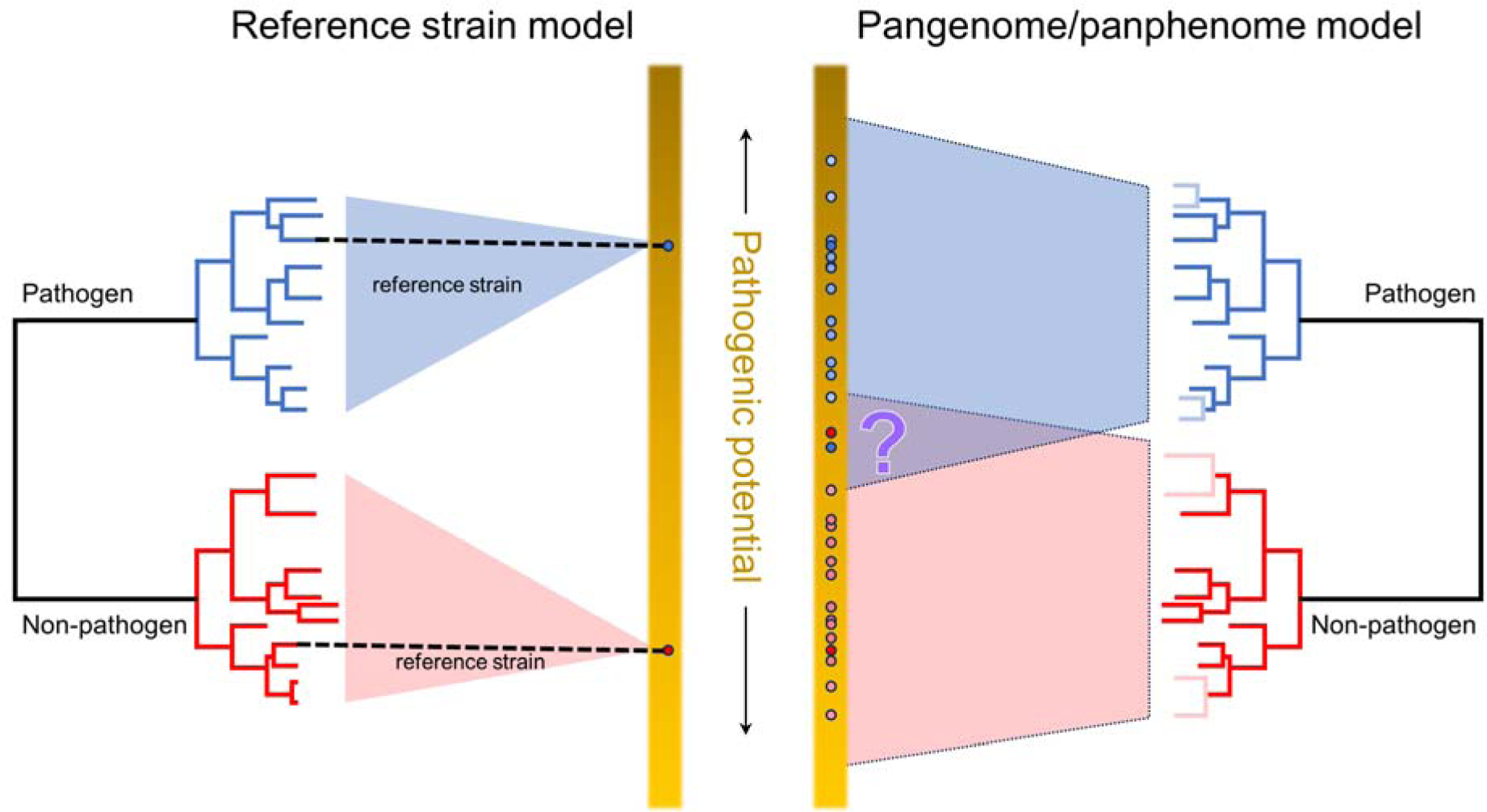
Generalized model of strain heterogeneity and its relevance for fungal pathogenicity. **Left**: Conventional, reference strain-based approach whereby all members of a fungal species are assigned the phenotypic features from one (or at most, a very few) reference strains. These reference strains may have been selected for their phenotypic extremes. This typological or essentialist approach views the phenotypic space of any species through the lens of the reference strain and thereby cannot accommodate population variation. Under this view, examination of the reference strains from two closely related species, one of which is a pathogen and one of which is non-pathogenic, reveals a large difference in their pathogenic potential. **Right**: A more comprehensive, population-level characterization of genetic and phenotypic traits across a diversity of strains. This approach stems from “population thinking” ^72^. In the context of pathogenicity, a population-thinking approach will increase emphasis of the reality of genetic and phenotypic variation exhibited by individual strains and can more fully capture the potential overlap (or its absence) in the pathogenic potential of closely related species.

Our results show that differences in pathogenic potential in *A. fischeri* cannot be adequately explained by genotype alone, and that they bear closer resemblance to observed differences in overall transcriptomic profiles under certain disease-relevant conditions (Figure 4A). These findings, coupled with the generally high prevalence of *A. fischeri* orthologs of genes previously identified as virulence-associated in *A. fumigatus* (Figure 4B), imply that virulence differences within *A. fischeri* are more likely the result of shifts in gene regulation than of any specific gene content differences.

In pathogenic species of *Aspergillus*, secondary metabolites comprise one of the better characterized sets of virulence factors that can have direct consequences on the host ^44, 45^. Each of these metabolites is the biochemical output of the protein products of specific BGCs, and one of the ways that metabolite levels can be altered is by the coordinated transcriptional regulation of the genes within the clusters. For example, our *A. fischeri* data show that most genes in the gliotoxin BGC are pervasively upregulated at 37°C in most strains, and gliotoxin itself is only detected at 37°C in several strains (Figure 5C).

Importantly, the observed strain heterogeneity in secondary metabolism within *A. fischeri* allowed us to identify a statistically significant association between variation in the production of just a few secondary metabolites and the breadth in pathogenic potentials between strains. Specifically, we observed a statistically significant association between strain virulence and the co-detection of gliotoxin and hexadehydroastechrome, raising the hypothesis that the production of just these two secondary metabolites in combination is sufficient for increasing a strain’s pathogenic potential (Figure S4). While we are not proposing that any specific secondary metabolite(s) should be credited with the broader gain/loss of a pathogenic phenotype within or across *Aspergillus* species, the strong effect size that can be attributed to the co-presence of just two metabolites offers support for the hypothesis that secondary metabolism-associated “cards” of virulence contribute to the “hand of cards”’ that enable any given *Aspergillus* strain or species to cause disease ^12, 20^.

The range of virulence observed between *A. fischeri* strains (Figure 2) suggests that a categorical classification of non-pathogen may not apply equally well to all *A. fischeri* strains and that reliance on single reference strains is unreliable. Rather than classifying all *A. fischeri* strains “non-pathogenic” based on the pathogenic potential of a single reference strain, our results argue that we should instead evaluate the pathogenic potential of a diversity of strains. Capturing strain-to-strain variation in virulence (and other infection-related genetic and phenotypic traits) would enable us to more accurately describe the pathogenic potential of entire species. Applied to enough strains of multiple species, we may begin to see convergences or even overlaps in pathogenic potentials between species (Figure 6), blurring the distinction of pathogens and non-pathogens.

If *A. fischeri* exhibits such a range of heterogeneity in virulence, what explains its apparent absence in the clinic? We offer two not mutually exclusive explanations: lack of ecological opportunity and species misidentification. *A. fischeri* occupies similar ecological niches in natural environments as its close relative and major pathogen, *A. fumigatus*; the two species frequently co-occur in decaying matter and soil across a range of environments and latitudes ^31, 59^. However, while *Aspergillus* species are frequently isolated from hospital environments ^60–62^, *A. fischeri* is less frequently present compared to more well-known *Aspergillus* pathogens, such as *A. fumigatus* or *A. flavus* ^63^. If correct, this general absence of *A. fischeri* in clinical spaces would deny it the same ecological opportunity enjoyed by other more abundant species.

The second possible explanation for the absence of *A. fischeri* in the clinic lies in the challenge of accurately identifying fungal species. On the one hand, with ∼150,000 species described from the likely millions of fungi inhabiting our planet ^64^, we are continuously discovering new species, including new pathogens such as the emerging pathogen *Candida auris* (described in 2009) and the cryptic pathogen *Aspergillus lentulus* (described in 2005) ^7, 65^. On the other hand, accurate classification of fungal species is nontrivial and requires molecular barcodes ^66, 67^. Because accurate, species-level identification of infectious agents is rarely a priority in clinical settings (particularly in cases of opportunistic fungal infections), cryptic species are often undetected.

Such imprecisions in identification can inject biases into the literature. For example, species identification based upon isolate morphology led to misattribution of several cases of invasive aspergillosis to *A. fumigatus,* but subsequent examination of molecular barcodes implicated another, morphologically indistinguishable, species in section *Fumigati* ^7^. Indeed, an in-depth, metagenomic survey of four clinical environments revealed not only the pervasive co-occurrence of *A. fumigatus* and *A. fischeri,* but the relative proportions of the two species remained remarkably constant across environments (approximately 7:1) ^63^. In contrast, clinical reports lean heavily on less-granular methods that are more expedient ^68^. Moving forward, we argue that a more comprehensive picture of the pathogenic potential of fungal species must include both the accurate taxonomic attribution of clinical isolates, their relatives, and the distinction within species between more- and less-virulent strains. Along these lines, aided by advances in genome sequencing technologies, calls for the adoption of “taxogenomic” approaches to species identification, especially in biomedically relevant clades of fungi that are increasing ^69–71^; implementation of such approaches would greatly increase our understanding of the species involved in fungal disease.

Variation in the pathogenic potentials of different strains then can be more deeply characterized, perhaps using moderate throughput *in vitro* assays (Figure 1), the results of which correspond well to the more-involved *in vivo* animal models of fungal disease (Figure 2). Metabolite profiling of these strains could then serve to examine how chemotypes evolve between/within species, possibly providing new insights into interplay between chemistry and pathogenic potential. Extension of this approach to strains in other species may reveal convergences in pathogenic potentials (Figure 6), providing not only evolutionary insights into how major pathogens originate, but also adding understanding to the underlying causes of virulence and effecting better clinical outcomes.

## METHODS

### *A. fischeri* strains analyzed in this study

All strains of *A. fischeri* were obtained from the culture collections of the Westerdijk Fungal Biodiversity Institute (Utrecht, The Netherlands). Strain identifiers and metadata are provided in Table 1.

### Genome sequencing and quality assessment

Each *A. fischeri* strain was grown at 30°C prior to DNA extraction. For long read sequencing, high molecular weight genomic DNA was isolated from pellets and sequenced by Oxford Nanopore (ONT) by SeqCenter (Pittsburgh, PA, USA). For short read DNA sequencing, genomic DNA was isolated using ZYMO RESEARCH Quick-DNA Fungal/Bacterial Kit, followed by TruSeq DNA PCR-Free library construction. Samples were then subject to 2 × 150 bp paired-end sequencing on an Illumina NovaSeq 6000 by VANTAGE NGS (Vanderbilt University, Nashville, TN, USA).

### *De novo* genome assembly

ONT reads from each strain were checked for contamination using Kaiju (v1.8.2) ^73^ and assembled using Flye (v.2.9-b1774) ^74^. Long-read assemblies were polished with the Illumina reads using *pilon* (v1.24) ^75^. The resulting assemblies were evaluated for completeness using BUSCO (fungi_odb10 and eurotiomycetes_obd10) ^76^. Assembly parameters were optimized for each strain so as to maximize BUSCO completeness. Generally, optimal assemblies were obtained by thresholding Flye for a read error of 0.02, followed by 2 rounds of pilon polishing.

Assemblies were evaluated for quality using QUAST (QUality ASsessment Tool; v.5.2.0) ^77^ and the best assembly (DTO15) was chosen to serve as a reference genome for all the other strains. Contigs from the DTO15 assembly were then scaffolded against the telomere-to-telomere assembly of *A. fumigatus* CEA10 ^78^, retaining the chromosome numbering of the CEA10 strain.

### Gene annotation

Gene model prediction and functional annotation were performed for each strain using Funannotate ^79^ (v.1.5.14). Briefly, assemblies were masked for repetitive elements using RepeatMasker ^80^. Transcript evidence for gene model training was obtained from the *de novo* assembly of RNAseq reads (see RNA sequencing section below) using Trinity ^81^. Gene models predicted to encode peptides shorter than 50 amino acids or transposable elements were removed. Transfer-RNA prediction was performed using tRNAscan-SE ^82^.

### Phylogenetics

To infer the strain phylogeny, single copy orthologs were extracted from the 16 *A. fischeri* genomes as well as two outgroup genomes of the *A. fumigatus* reference strains Af293 ^83^ and CEA10 ^78^ using BUSCO (v. 5.4.6) ^76^ and the eurotiomycetes_odb10 data base (N=3,546 BUSCO entries). After filtering for BUSCO genes present across all strains and the outgroups (N=3,375, >96%), the remaining loci were aligned with Muscle5 ^84^. Manual inspection and curation identified 21 poorly identified and/or badly aligning sequences that were discarded. This process resulted in final multiple sequence alignments containing 3,354 genes, which were concatenated and used to build the strain phylogeny with IQ-TREE (v. 2.2.2.3) ^85^.

### Pangenome analysis

The *A. fischeri* pangenome was constructed using Pantools (v4.1.1) ^86, 87^. In short, a database was created using the amino acid sequences for each of the gene models produced by Funnanotate for each of the *A. fischeri* strains. Initial orthogroups were determined using the Pantools’ `build_panproteome` function, then the `optimal_`grouping function identified the optimal sequence similarity threshold (75%) for maximizing precision and recall (the F-score). Initial assessment of core, accessory, and singleton status of orthogroups was performed using the `gene_classification` function with default settings. The `gene_classification` function also provided a Heap’s Law calculation.

Next, orthogroups from Pantools were clustered by sequence similarity with a dynamic similarity threshold using ALFATClust (version available as of 04-10-2023) with default settings (Chiu & Ong 2022). This resulted in a set of 16,148 (16,427 including 279 multi-copy genes) gene clusters across all 16 strains. Each of these gene clusters were assigned a number within the parent orthogroup. Gene clusters were characterized as core, accessory, or singletons based on the number of strains represented in each cluster. Core genes were those gene clusters that contained sequences from all 16 strains, accessory genes contained sequences from 2-15 strains, and genes found in a single strain were considered singletons. Gene copy number was assessed by counting the number of sequences from each strain within a gene cluster. In downstream analysis, multicopy gene clusters (gene clusters that contained multiple genes from a single strain) and gene clusters whose average percent identity was lower than 70% were excluded from analysis.

To relate *A. fischeri* pangenes back to the canonical reference strains of *A. fumigatus* (Af293) and *A. fischeri* (NRRL 181), we sourced annotations for Af293 and NRRL181 from FungiDB release 62 ^88^. These reference sequences were used to construct a custom BLAST database against which, pangenecluster centroids (identified using DIAMOND (v2.1.6)^89^) were reciprocally BLASTed Any duplicate BLAST results were resolved by selecting for maximum bit score. The homologous genes found were used for annotating the pangenome.

### Population genetics

Population genetic analyses were performed with GATK (v. 4.1.2.0-Java-1.8.0_192) ^90^. DTO15 was designated the reference genome, which was indexed using SAMtools (v. 1.16.1) ^91^, BWA (v 0.7.17) ^92^, and GATK. To produce a high-confidence SNV set (known_sites file), we initially ran GATK in default diploid mode applying quality filtering with VCFtools (v. 0.1.16) ^93^ and the following flags: --minGQ 20, --minDP 5, -min-meanDP 10, --mac 3 to the preliminary population VCF file. We extracted the resulting genotype table and used sed to remove all heterozygous SNVs. This set of high-confidence SNVs became the known_sites file. We then performed the analysis following the “Germline short variant discovery (SNVs + Indels)” best practices (https://gatk.broadinstitute.org/hc/en-us/articles/360035535932-Germline-short-variant-discovery-SNVs-Indels-). Duplicates were marked, base quality score recalibration (BQSR) was performed with the custom known_sites file, and SNVs were called with GATK’s HaplotypeCallerSpark, setting sample-ploidy to 1; a total of 303,690 SNVs were identified. Applying the traditional/alternative hard filtering ^94^ did not remove any SNVs.

To infer population structure, we used ADMIXTURE (v1.3.0) ^95^ on the set of 303,690 SNVs. These SNVs were first filtered on allele frequency using *vcftools* ^93^ (resulting in 215,627 SNVs with an allele frequency greater than 10%), LD pruned in PLINK ^96^, then clustered by PCA to create input for ADMIXTURE. Finally, ADMIXTURE was run using both 5- and 10-fold cross validation and K values were selected based upon local minima of cross validation errors.

### Biosynthetic Gene Cluster detection

To identify biosynthetic gene clusters (BGCs), we collected the genes and locus tags from primary literature and the BGC repository MIBiG (v3.1) ^97^. BGCs in the *A. fischeri* strains were annotated by antiSMASH (v6.0.1) ^98^ and manually reviewed to confirm gene content, borders and syntenic similarity.

### RNAseq sample preparation and RNA isolation

For RNAseq preparation, conidial suspensions from 15 strains were harvested from 3 days old-conidia cultured on complete media (strain DTO7 was excluded because of poor growth at 37°C). The conidial suspension was counted using a hemocytometer and adjusted to a final concentration of 1×10^8^ conidia/ml. For the two experimental conditions, conidia were cultured in 250 mL flasks containing 30 mL minimal media and incubated at either 30°C or 37°C for 24 hours with agitation.

For RNA extraction, material was pelleted by centrifugation, the pellet separated by filtration, rinsed with water, flash frozen, and stored at −80°C. For total RNA extraction, the mycelial biomass (around 100 mg of tissue) was first homogenized by grinding in liquid nitrogen and then RNA was isolated using TRIzol (Invitrogen, Catalog Number 15596026) and cleaned up with the RNeasy clean-up method (Qiagen, Catalog Number 74104). Finally, the RNA extract was treated using DNase (Qiagen cat#79254) and was quantified using a spectrophotometer. The purified RNA was stored at −80°C.

### RNA sequencing

RNA samples submitted to Vanderbilt’s sequencing center VANTAGE NGS (Vanderbilt University, Nashville, TN, USA), where RNA was again checked for RNA integrity poly-A enriched prior to Illumina preparation using NEBNext® Poly(A) selection followed by paired-end sequencing (150bp from each end) on an Illumina NovaSeq 6000 S4.

### RNAseq read alignment and differential expression

Reads were quality checked using FastQC (v0.11.9)^99^ and filtered using Trimmomatic (v0.040-rc1)^100^. Reads from each strain were aligned back to that strain’s genome assembly and unique set of gene models using STAR (v2.7.10b) ^101^. Unique read counts per strain were then gathered for each sample using HTseq (v.2.0.3) ^102^. Counts were then evaluated on a strain-by-strain basis for differential expression between the 30°C and 37°C treatment groups (three replicates per treatment using DEseq2 (v1.42.0) ^103^. DEseq2 gene names were standardized across all samples using the pangenome results. Genes were considered to be significantly differentially expressed if their false discovery rate-corrected (Benjamini and Hochberg method) p-value (padj) was less than 0.05.

### Growth and extraction of *Aspergillus fischeri* cultures for chemical analyses

*A. fischeri* strains (n=16) were grown in oatmeal cereal media (Old fashioned breakfast Quaker oats) for chemical characterization ^29^. Briefly, fungal cultures were started by aseptically cutting mycelia from a culture growing in a Petri dish on malt extract agar (MEA, Difco), and this material was transferred to a sterile Falcon tube with 10 mL of liquid YESD media (YESD; 20 g soy peptone, 20 g dextrose, 5 g yeast extract, 1 L distilled H2O). This seed culture was grown for 7 days on an orbital shaker (∼100 rpm) at room temperature (∼23°C) and used to inoculate the oatmeal cereal media, which was prepared by adding 10 g of oatmeal in a 250 mL, wide mouth Erlenmeyer flask with ∼17 mL of deionized water followed by autoclaving at 121°C for 30 min. For each of the 16 strains, three flasks of each of the samples (i.e., three biological replicates per sample) were incubated for 10 d at 30°C and three at 37°C. Since it is known that light modulates growth, sexual reproduction, and secondary metabolite production in fungi ^104^, the incubators were equipped with LED lamps (Ustellar, flexible LED strip lights; 24 W) with light regulated into 12:12 h dark/light cycles ^105^.

At the end of 10 d, each individual flask was extracted by adding 60 mL of CHCl_3_-MeOH (1:1), chopped with a spatula, and shaken overnight (∼16 h) at ∼100 rpm at room temperature. The cultures were then filtered *in vacuo*, and 90 mL of CHCl_3_ and 150 mL of deionized water were added to each of the filtrates. The mixture was then transferred to a separatory funnel and shaken vigorously. The organic layer *(i.e*., bottom layer) was drawn off and evaporated to dryness *in vacuo*. The dried organic layer was reconstituted in a 100 mL of CH_3_CN-MeOH (1:1) and 100 mL of hexanes, transferred to a separatory funnel, and shaken vigorously. The defatted organic layers were evaporated to dryness.

All the defatted organic layers were analyzed individually by UPLC-HRESI-MS utilizing a Thermo LTQ Orbitrap XL mass spectrometer equipped with an electrospray ionization source. A Waters Acquity UPLC was utilized with a BEH C18 column (1.7 μm; 50 mm × 2.1 mm) set to 40°C and a flow rate of 0.3 mL/min. The mobile phase consisted of a linear gradient of CH_3_CN−H_2_O (both acidified with 0.1% formic acid), starting at 15% CH_3_CN and increasing linearly to 100% CH_3_CN over 8 min, with a 1.5 min hold before returning to the starting conditions. The profile of secondary metabolites of all the extracts was examined using well-established procedures ^29^.

### Isolation and identification of secondary metabolites

After comparison of the UPLC-HRESI-MS data for each biological replicate, the defatted organic layers for each strain and condition were combined due to the similarity of their chemical profiles, so as to generate a larger pool of material for isolation studies, which were carried out using well-established natural products chemistry procedures ^106, 107^. The extracts were first fractionated by normal phase flash chromatography, and the fungal metabolites were isolated using preparative HPLC.

The isolated fungal metabolites were identified by direct comparison of the spectroscopic and spectrometric properties with those reported previously, and where possible, structures were validated by comparisons with authentic reference standards ^107^. Additionally, mass defect filtering^108^ was used to identify structurally-related analogues of the isolated compounds.

The NMR data were collected using a JEOL ECS-400 spectrometer operating at 400 MHz for ^1^H and 100 MHz for ^13^C, an JEOL ECA-500 spectrometer operating at 500 MHz for ^1^H and 125 MHz for ^13^C or an Agilent 700 MHz spectrometer, equipped with a cryoprobe, operating at 700 MHz for ^1^H and 175 MHz for ^13^C. The HPLC separations were performed on a Varian Prostar HPLC system equipped with a Prostar 210 pump and a Prostar 335 photodiode array detector (PDA), with the collection and analysis of data using Galaxy Chromatography Workstation software. The columns used for separations were either a Synergi C18 preparative (4 μm; 21.2 × 250 mm) column at a flow rate of 21.2 mL/min, Luna PFP(2) preparative (5 μm; 21.2 × 250 mm) column at a flow rate of 17 mL/min or an Gemini NX-C18 preparative (5 μm; 19 × 250 mm) column at a flow rate of 17 mL/min. Flash chromatography was performed on a Teledyne ISCO Combiflash Rf 200 and monitored by ELSD and PDA detectors.

### Principal component analysis of metabolomics results

Principal component analysis (PCA) and hierarchical clustering were performed on the UPLC-HRESI-MS data. Untargeted UPLC-HRESI-MS datasets for each sample were individually aligned, filtered, and analyzed using Mzmine (v2.53) (https://sourceforge.net/projects/mzmine/). Peak list filtering and retention time alignment algorithms were used to refine peak detection, and the join algorithm integrated all sample profiles into a data matrix using the following parameters: Mass detection: MS1 positive mode. ADAP: Group intensity threshold: 20000, Min highest intensity: 60000, m/z tolerance: 0.003. Chromatogram deconvolution: Wavelets (ADAP) algorithm. Join aligner and gap filling: retention time tolerance: 0.05 min, m/z tolerance: 0.0015 m/z. The resulting data matrix was exported to Excel (Microsoft) for analysis as a set of m/z–RT (retention time) pairs with individual peak areas. Samples that did not possess detectable quantities of a given marker ion were assigned a peak area of zero to maintain the same number of variables for all sample sets. Ions that did not elute between 1 and 10 min and/or had an m/z ratio < 200 or > 900 Da were removed from analysis. Relative standard deviation was used to understand the quantity of variance between the injections, which may differ slightly based on instrument variance. A cutoff of 1.0 was used at any given m/z–RT pair across the biological replicate injections, and if the variance was greater than the cutoff, it was assigned a peak area of zero. PCA analysis and hierarchical clustering were created with Python. The PCA scores plots were generated using the averaged data of the three individual biological replicates.

### Murine model of pulmonary aspergillosis

Wild-type BALB/c female mice, body weight 20 to 22 g, aged 8-9 weeks, were kept in the Animal Facility of the Laboratory of Molecular Biology of the School of Pharmaceutical Sciences of Ribeirão Preto, University of São Paulo (FCFRP/USP), in a clean and silent environment, under normal conditions of humidity and temperature, and with a 12:12 h dark/light cycle. The mice were given food and water ad libitum throughout the experiments. Mice were immunosuppressed with cyclophosphamide (150 mg per kg of body weight), which was administered intraperitoneally on days −4, −1, and 2 prior to and post infection. Hydrocortisonacetate (200mg/ kg body weight) was injected subcutaneously on day −3. *A. fischeri* strains and the *A. fumigatus* CEA17 strain, a clinically derived and highly virulent strain, that we used as a positive control, were grown on minimal media for 2 days prior to infection. Fresh conidia were harvested in PBS and filtered through a Miracloth (Calbiochem). Conidial suspensions were spun for 5 min at 3,000 x g, washed three times with PBS, counted using a hemocytometer, and resuspended at a concentration of 5.0 x 106 conidia/ ml. The viability of the administered inoculum was determined by incubating a serial dilution of the conidia on minimal media at 37°C. Mice were anesthetized by halothane inhalation and infected by intranasal instillation of 1.0 x 105 conidia in 20 µl of PBS. As a negative control, a group of 10 mice received PBS only. Mice were weighed every 24 h from the day of infection and visually inspected twice daily. The statistical significance of comparative survival values was calculated by Prism statistical analysis package by using the Log-rank (Mantel-Cox) Test and Gehan-Breslow-Wilcoxon tests.

The virulence of each strain of *A. fischeri* was evaluated in cohorts of 20 mice (10 treatments, 10 negative PBS controls). A positive control cohort of 20 mice was also evaluated (10 *A. fumigatus* CEA17 treatments and 10 negative PBS controls). Two replicates of each cohort were conducted.

The principles that guide our studies are based on the Declaration of Animal Rights ratified by UNESCO on January 27, 1978 in its 8th and 14th articles. All protocols adopted in this study were approved by the local ethics committee for animal experiments from the University of São Paulo, Campus of Ribeirão Preto (Permit Number: 08.1.1277.53.6; Studies on the interaction of *A. fischeri* with animals). Groups of five animals were housed in individually ventilated cages and were cared for in strict accordance with the principles outlined by the Brazilian College of Animal Experimentation (COBEA) and Guiding Principles for Research Involving Animals and Human Beings, American Physiological Society. All efforts were made to minimize suffering. Animals were clinically monitored at least twice daily and humanely sacrificed if moribund (defined by lethargy, dyspnea, hypothermia, and weight loss). All stressed animals were sacrificed by cervical dislocation.

### Macrophage cytokine response to conidia

*Aspergillus* strains were cultivated on minimal medium agar plates at 37°C for 3 days. Conidia were harvested in sterile water with 0.05 % (vol/vol) Tween20. The resulting suspension was filtered through two layers of gauze (Miracloth, Calbiochem). The conidial concentration was determined using a hemocytometer.

BALB/c bone marrow-derived macrophages (BMDMs) were obtained as previously described ^109^. Briefly, bone marrow cells were cultured for 7–9 days in RPMI 20/30, which consists of RPMI-1640 medium (Gibco, Thermo Fisher Scientific Inc.), supplemented with 20 % (vol/ vol) FBS and 30 % (vol/vol) L-Cell Conditioned Media (LCCM) as a source of macrophage colony-stimulating factor (M-CSF) on non-treated Petri dishes (Optilux - Costar, Corning Inc. Corning, NY). Twenty-four hours before experiments, BMDM monolayers were detached using cold phosphate-buffered saline (PBS) (Hyclone, GE Healthcare Inc. South Logan, UT) and cultured, as specified, in RPMI-1640 (Gibco, Thermo Fisher Scientific Inc.) supplemented with 10 % (vol/vol) FBS, 10 U/mL penicillin, and 10 mg/mL streptomycin, (2 mM) L-glutamine, (25 mM) HEPES, pH 7.2 (Gibco, Thermo Fisher Scientific Inc.) at 37°C in 5 % (vol/vol) CO_2_ for the indicated periods.

BMDMs were seeded at a density of 10^6^ cells/ml in 24-well plates (Greiner Bio-One, Kremsmünster, Austria). The cells were challenged with the conidia of different strains at a multiplicity of infection of 1:10 and incubated at 37°C with 5 % (vol/vol) CO_2_ for 48h. BMDMs were also stimulated with lipopolysaccharide (LPS; standard LPS, E. coli 0111: B4; Sigma-Aldrich, 500 ng/mL) plus Nigericin (tlrl-nig, InvivoGen 5 μM/mL) and cell medium alone, which were used respectively as the positive and negative controls. Cell culture supernatants were collected and stored at −80°C until they were assayed for TNF-α, IL-1, and IL-6 release using Mouse DuoSet ELISA kits (R&D Systems, Minneapolis, MN, USA, according to the manufacturer’s instructions. For cytokine determination, plates were analyzed by using a microplate reader (Synergy™ HTX Multi-Mode, BioTek) measuring absorbance at 450 nm. Cytokine concentrations were interpolated from a standard curve and statistical significance was determined using an ANOVA.

### Macrophage killing assay

BMDMs were seeded at a density of 10^6^ cells/ml in 24-well plates (Corning^®^ Costar^®^) and were challenged with conidia at a multiplicity of infection of 1:10 and incubated a 37°C with 5 % (vol/vol) CO_2_ for 24h. After incubation media was removed the cells were washed with ice-cold PBS and finally 2 ml of sterile water was added to the wells. A P1000 tip was then used to scrape away the cell monolayer and the cell suspension was collected. This suspension was then diluted 1:1000 and 100 μl was plated on Sabouraud agar before the plates were incubated at 37°C overnight and the colonies were counted. 50 μl of the inoculum adjusted to 10^3^/ml was also plated on SAB agar to correct CFU counts. The CFU/ml for each sample was calculated and compared to the A1160 wild-type strain using (GraphPad Prism 8.0, La Jolla, CA). All assays were performed in four replicates in two independent experiments.

### Growth phenotyping

All strains were cultured from conidial glycerol stock (stored at −80°C) on Complete Minimal media agar plates (20 g of D-glucose, 50 mL of 20x salt solution [120 g of NaNO_3_, 10.4 g of KCI, 10.4 g of MgSO_4_ x 7H20 and 30.4 g of KH_2_PO_4_ g/L], 1 mL Trace Element Solution [22 g of ZnSO_4_ x7H_2_O, 11 g of H_3_BO_3_, 5 g of MnCl_2_x4H_2_O, 5 g of FeSO_4_x7H_2_O, 1.6 g of COCl_2_x 5H_2_O, 1.6 g of CuSO_4_ x5H_2_O, 1.1 g of (NH_4_)_6_Mo_7_O_24_x4H_2_O, and 50 g of Na_4_EDTA], 1 g of Yeast Extract, 2 g of Peptone, and 1.5 g of Casamino Acid g/L, 1.5% of agar, pH adjusted to 6.5) at 30°C under dark conditions in a culture chamber (Thermo Fisher, MaxQ6000). A 6 mm cork borer was used to obtain edge mycelia and subculture on Minimal Media Agar (50 mL 20x solution, 1 mL Trace Solution, and 10 g of D-glucose, 20% of agar and adjusted pH at 6.5) at 30°C and 37°C under dark conditions. After 2 days of growth, the radial growth of the mycelial mats for each strain was quantified manually using a ruler. Three replicates were done per strain, per temperature condition.

## DATA AVAILABILITY

Sequencing data and genome assemblies associated with this project will become available through GenBank upon publication. The NMR data will be deposited in the Natural Products Magnetic Resonance Database (https://npmrd-project.org/). All other data necessary to replicate this work will be available on FigShare.

## ACKNOWLEDGMENTS

We thank members of the Rokas lab for helpful discussions and feedback. Computational infrastructure at Vanderbilt University was provided by The Advanced Computing Center for Research and Education (ACCRE). This research was supported by the National Institutes of Health National Institute of Allergy and Infectious Diseases (R01 AI153356). The content is solely the responsibility of the authors and does not necessarily represent the official views of the National Institutes of Health. Research in A.R.’s lab is also supported by the National Science Foundation (DEB-2110404) and the Burroughs Wellcome Fund. Research in G.H.G.’s lab is also supported by the Fundação de Amparo à Pesquisa do Estado de São Paulo (FAPESP) grant numbers 2021/04977-5 (G.H.G.) and 2023/00206-0 (ED), the Conselho Nacional de Desenvolvimento Científico e Tecnológico (CNPq) grant numbers 301058/2019-9 and 404735/2018-5 (G.H.G.), and the CNPq, FAPESP and Fundação Coordenação de Aperfeiçoamento do Pessoal do Ensino Superior (CAPES) grant number 405934/2022-0 to G.H.G. (The National Institute of Science and Technology INCT Funvir), all from Brazil. G.H.G.’s lab was also funded by the Joint Canada-Israel Health Research Program, jointly supported by the Azrieli Foundation, Canada’s International Development Research Centre, Canadian Institutes of Health Research, and the Israel Science Foundation. This work was performed in part at the Joint School of Nanoscience and Nanoengineering, a member of the National Nanotechnology Coordinated Infrastructure (NNCI), which is supported by the National Science Foundation (NSF; Grant ECCS-2025462).

## CONFLICT OF INTEREST

A. R. is a scientific consultant for LifeMine Therapeutics, Inc.

