## Supplementary Figures for "Strain heterogeneity in a non-pathogenic fungus highlights factors contributing to virulence"

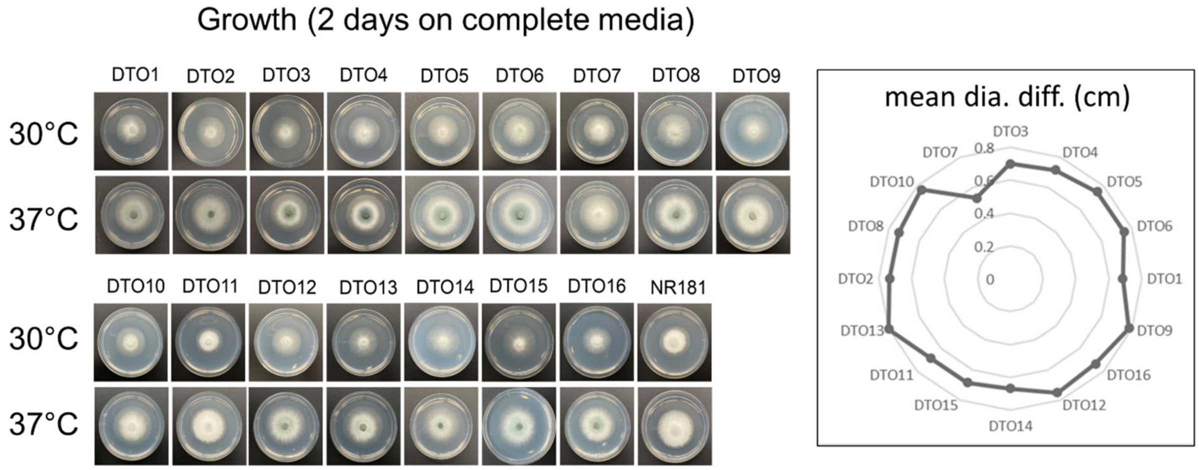


**Figure S1: Growth assay reveals differences between strains and temperatures**.
**Left**: Representative images of the 16 *A. fischeri* strains used in this study following 2 days of growth on complete minimal media at 30°C and 37°C. **Right**: Radar chart shows the amount of additional growth diameter observed at 37°C compared to 30°C (the mean differences in size of 3 replicates of each strain are shown). At 37°C, growth was substantially accelerated in all strains, with most strains displaying some degree of conidial melanization that was not present at 30°C. The sole exception was DTO7, which grew much more slowly.

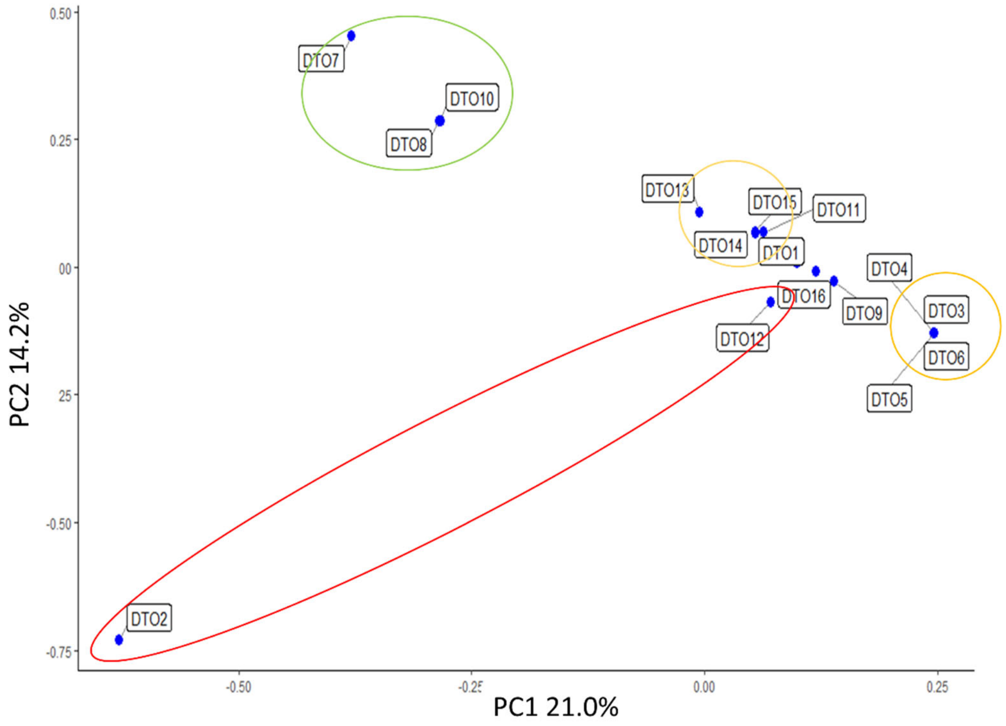


**Figure S2: Single nucleotide variant-based distances consistent with intraspecific relationships inferred by other methods**. Principal components analysis (PCA) of intraspecific distances from the subset of single nucleotide variants (SNVs) used in the population assignment analysis using the ADMIXTURE package. PCA distances between strains are consistent with relationships revealed by genetic structure analyses using DAPC (Discriminant Analysis of Principal Components) (Figure 3C) and the strain phylogeny (Figure 3A).

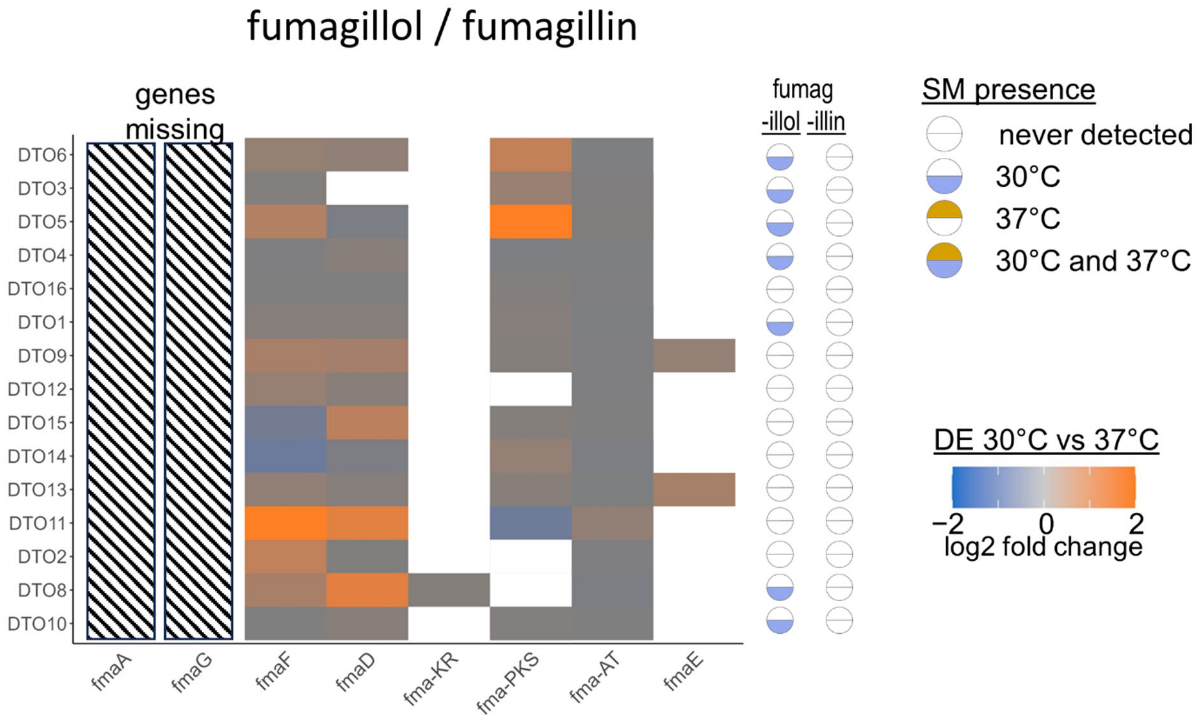


**Figure S3: Fumagillin BGC in all *A. fischeri* genomes lacks genes present in the canonical version yet intermediate products are detectable.**
Temperature-dependent differential expression of genes of the fumagillin BGC. Orthologs of the *fmaA* (*Afu8g00520*) and *fmaG* (*Afu8g00510*), both of which are present in the canonical version of the *A. fumigatus* BGC, are absent in the genomes of all *A. fischeri* strains (hatched areas). However, fumagillol, an intermediate metabolite of the fumagillin biosynthetic pathway, is detected in seven strains at 30°C. This suggests that the *A. fischeri* pathway for fumagillin biosynthesis is at least partially functional in some strains.

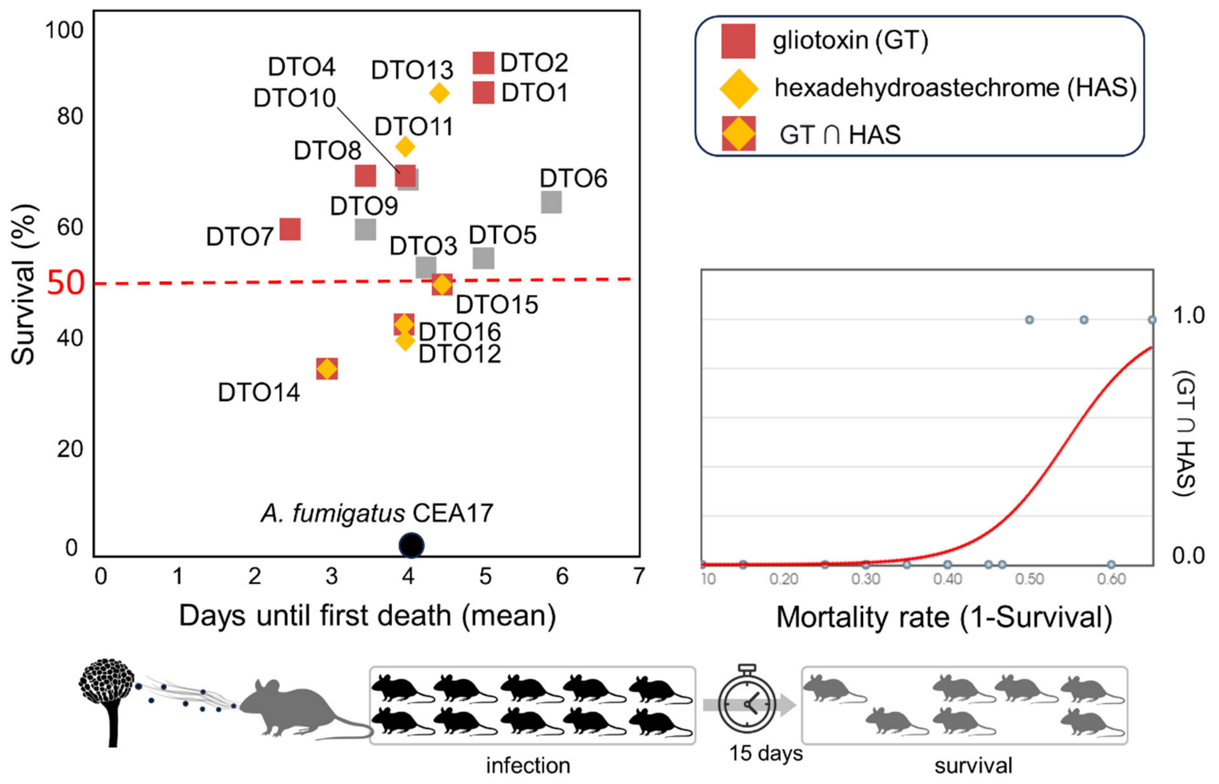


**Figure S4: Production of both the gliotoxin and hexadehydroastechrome secondary metabolites is significantly associated with *A. fischeri* virulence.**

**Left**: Occurrence of the gliotoxin and hexadehydroastechrome secondary metabolites with respect to virulence in mice (*c.f.*, Figure 2). Both secondary metabolites are detected at 37°C in 3 of the 4 most virulent strains (DTO14, DTO15, and DTO16; ≥ 50% mortality). **Right**: Logistic regression curve (red line) where the independent variable (x-axis) is mortality in a mouse model of pulmonary aspergillosis, and the dependent variable (y-axis) is indicating the presence (1) or absence (0) of the [gliotoxin + hexadehydroastechrome] metabolite pair. This model provides support to the hypothesis that higher mortality can be confidently related back to the simultaneous presence of these compounds in *A. fischeri* strains ($\chi^2$=7.9913, p value=0.0047).
